# Cross-state representation learning enables prospective discovery of dynamic protein allosteric sites

**DOI:** 10.64898/2026.11.09.765238

**Authors:** Qiuhan Ren, Liying Wu, Jinyin Zha, Mingyu Li, Wanting Feng, Haijiao Zhang, He Yang, Rui Li, Xinyi Liu, Jixiao He, Wanyao Zhou, Yuchengze Song, Ruidi Zhu, Chengwei Wu, Qiancheng Shen, Xi Cheng, Wenming Qin, Zhen Zheng, Jian Zhang

**Author notes:** Corresponding authors (Jian Zhang); (Xi Cheng); (Wenming Qin).

## Abstract

Allosteric regulation offers multifunctional induction in therapeutic interventions, yet conformational plasticity hinders computational identification of allosteric sites. Machine learning models rely on static *holo* (ligand-bound) structures, generalizing poorly to *apo* or substrate-bound states where binding pockets are transient or cryptic. We present AsBench2, a cross-state site-recognition benchmark, and AllositeUltra, a deep learning framework integrating multimodal protein representations (combining structure-aware language models with physical dynamics) and residue-guided candidate supplementation, effectively decoupling site detection from static pocket geometry. AllositeUltra demonstrates superior generalization, achieving Top-3 site recovery rates of 65.2% and 70.6% on *apo* and substrate-bound structures, respectively, representing relative improvements of 36.4% and approximately 50% over the best-performing baselines. Prospective application to MAGI2 identifies a previously unrecognized interdomain allosteric pocket, validated at atomic resolution through small-molecule screening and X-ray crystallography. Proteome-wide application yields a high-confidence atlas of targetable allosteric sites, bridging the gap between structural representation learning and prospective AI-driven drug discovery.

## Introduction

Allosteric regulation enables small molecules to modulate protein function by binding outside orthosteric sites, altering conformational populations and dynamic ensembles^1,2^. This mechanism offers unique opportunities to achieve exquisite target selectivity^3,4^, activate enzymes^5^, and disrupt traditionally “undruggable” targets^6^. Consequently, structure-based allosteric drug discovery^7^ relies fundamentally on the accurate computational identification of potential allosteric sites^8^.

However, identifying allosteric sites remains a formidable challenge for structure-based computational models. In prospective drug discovery, target structures are predominantly available only in unliganded (*apo*) or substrate-bound states. In these conformations, dynamic local rearrangements often contract, distort, or occlude pockets that are fully exposed only upon allosteric ligand binding^9–11^. Existing deep learning and geometric representation algorithms heavily depend on explicitly preformed, static pocket geometries^12–14^. When local backbone or side-chain fluctuations alter cavity shapes, these models frequently fail to generate valid candidate regions or recognize corresponding regulatory sites across different structural states. Furthermore, current computational benchmarks are overwhelmingly dominated by allosteric ligand-bound (*holo*) structures, creating a substantial evaluation bias that overestimates real-world prospective performance and masks the cross-state generalization bottleneck.

Current computational methods leverage pocket geometry, local physicochemical properties^12,13^, protein dynamics^15–17^, and sequence embeddings^14^ to identify candidates. Yet, because conformational transitions reconfigure local contact networks and feature representations, cues optimized on *holo* structures degrade significantly when applied to alternative states. While curated databases such as ASD^18–22^ and AsBench^23^ have advanced allosteric research, a standardized benchmark designed specifically to test cross-state recognition across structural states has been lacking.

To address these challenges, we establish AsBench2, a curated, cross-state benchmark derived from experimentally validated allosteric sites linked across *holo*, *apo*, and substrate-bound structures. Building on this framework, we develop AllositeUltra, a novel multimodal deep learning architecture designed to decouple allosteric site discovery from static cavity geometry. AllositeUltra fuses structure-aware protein language model representations, local physicochemical attributes, dynamic response profiles, and protein-wide inter-residue attention networks. Crucially, it introduces a residue-guided candidate supplementation mechanism that generates and evaluates candidate regions directly from learned allosteric residue probabilities, allowing the model to uncover cryptic or closed sites missed by traditional geometric detection algorithms.

We demonstrate that AllositeUltra significantly outperforms existing state-of-the-art baselines in recovering allosteric sites from *apo* and substrate-bound structures. We validate the functional relevance of predicted novel pockets through site-directed mutagenesis across diverse protein families, and show that AllositeUltra predictions effectively guide downstream ligand docking and pose prediction workflows. In a prospective application against MAGI2, model-guided screening successfully identifies a micromolar ligand targeting a previously unrecognized interdomain site, which we confirm at 2.24 Å resolution via X-ray crystallography. Finally, applying AllositeUltra proteome-wide yields a comprehensive atlas of human candidate allosteric sites. Together, AsBench2 and AllositeUltra establish a powerful framework for cross-state structural learning, expanding the targetable space for prospective allosteric drug discovery.

## Results

### Existing ligand-bound benchmarks overestimate prospective allosteric-site recovery

To test whether recognition performance in allosteric ligand-bound structures extends to conformations available during discovery, we collected additional *apo* and substrate-bound structures corresponding to known sites in AsBench^23^ and assembled a paired challenge set (Supplementary Table 1). We evaluated three commonly used methods with distinct modelling strategies—DeepAllo, AllositePro and PASSERRANK^13,14,24^—on allosteric ligand-bound structures and their *apo* or substrate-bound counterparts. Relative to the allosteric ligand-bound state, Top-3 success rates decreased by 26.3–93.1% in *apo* structures and 70.8–94.3% in substrate-bound structures (Fig. 1c). Top-1 success rates and recall showed similar declines (Supplementary Fig. 1).

**Fig 1.**
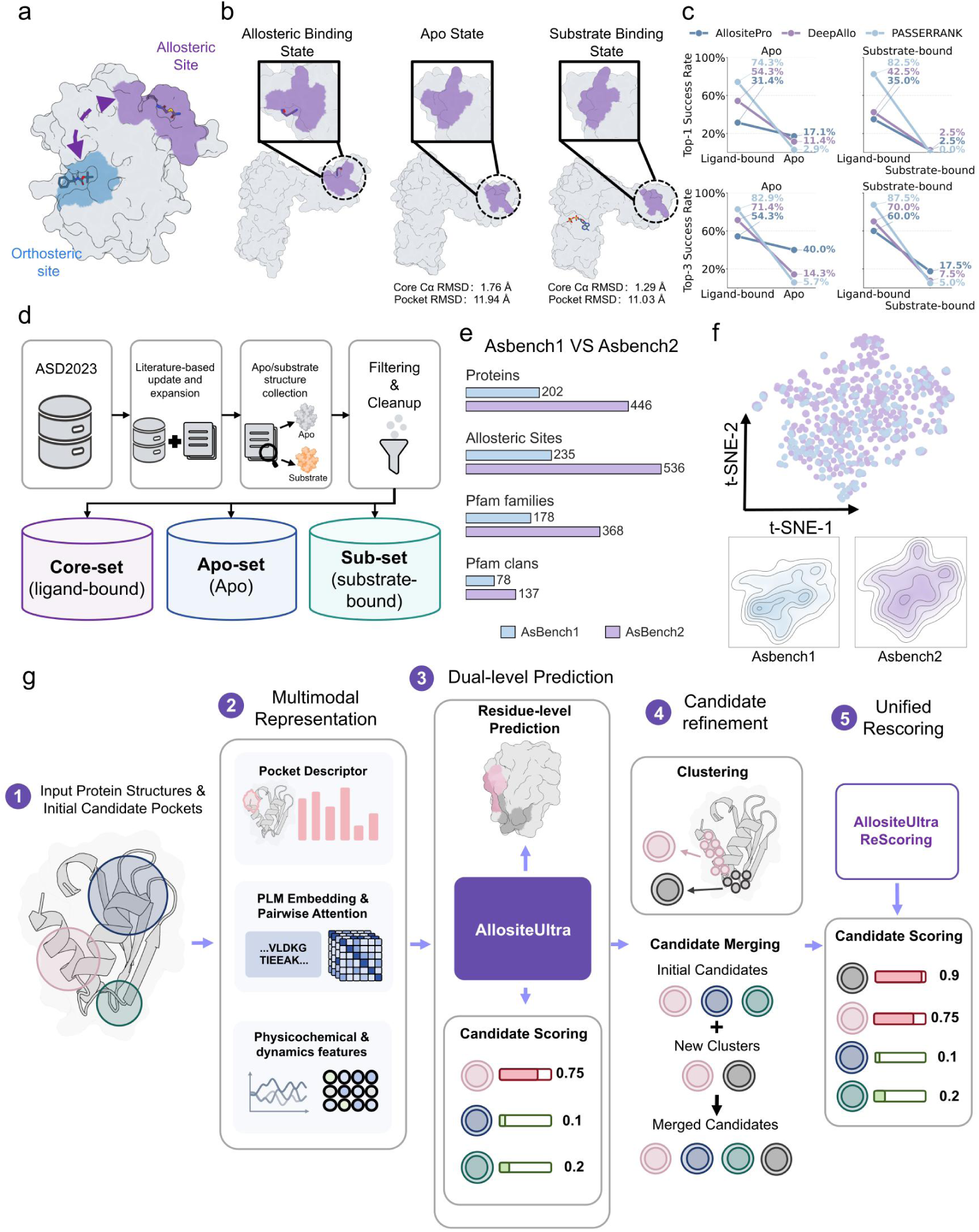
AsBench2 construction and cross-state allosteric-site prediction with AllositeUltra. a, Schematic of the spatial relationship between allosteric and orthosteric sites. b, Structures and pocket geometries of Mycobacterium tuberculosis ATP phosphoribosyltransferase bound to histidine and AMP (PDB: 1NH8), in the *apo* state (PDB: 1NH7) ^25^, and bound to ATP (PDB: 5U99) ^26^. Core Cα RMSD and pocket RMSD denote root-mean-square deviations of protein core Cα atoms and pocket residues, respectively. c, Top-1 and Top-3 success rates of DeepAllo, AllositePro and PASSERRANK for recovering the same allosteric sites in allosteric ligand-bound structures and their corresponding *apo* or substrate-bound structures. d, AsBench2 construction workflow. e, Comparison of dataset size and biological coverage between AsBench1 ^23^ and AsBench2. f, t-SNE visualization of protein embeddings from AsBench1 and AsBench2. g, AllositeUltra workflow.

Even when the analysis was extended to all candidate pockets generated by each method, target-site coverage still differed markedly across structural states. Across methods, 0–13.6% of allosteric ligand-bound systems lacked any candidate satisfying the geometric site-matching criteria. The corresponding proportions increased to 45.7–80.0% for *apo* systems and 45.0–90.0% for substrate-bound systems (Supplementary Table 2). Some sites were thus missed during candidate generation, limiting subsequent recovery through scoring and ranking. Even when changes in overall protein structure are limited, local contraction or rearrangement may alter pocket shape and physicochemical environment, making the same site difficult to recognize in another structural state (Fig. 1b). Benchmarks based on allosteric ligand-bound structures therefore do not adequately capture the difficulty of prospective allosteric-site discovery, motivating a new benchmark for systematically evaluating recovery of the same sites across structural states.

### AsBench2 enables systematic cross-state evaluation of allosteric sites

We therefore constructed AsBench2 to enable systematic evaluation of cross-state recognition. Starting from ASD2023^22^, we updated and filtered allosteric-site records using evidence from functional experiments in primary studies and ligand-bound crystal structures. Sequence and structural alignments linked these sites to corresponding *apo* and substrate-bound structures, establishing correspondence for the same experimentally supported sites across conformations (Fig. 1d). The Core-set comprises 536 allosteric sites across 446 proteins. The corresponding Apo-set and Sub-set comprise 387 *apo* (unliganded) and 275 substrate-bound structures, respectively, with the target allosteric site unoccupied in both sets.

Relative to AsBench1^23^, AsBench2 contains approximately 2.3 times as many site records and covers 2.1 times as many Pfam families^27^ and 1.8 times as many clans (Fig. 1e). This expansion spans multiple species and encompasses major enzyme classes alongside membrane, transport, signalling and transcription-related proteins. A t-SNE visualization of protein language-model representations showed a broader distribution for AsBench2 than for AsBench1 (Fig. 1f). By combining consistent experimental site annotations with structural correspondence across states, AsBench2 enables systematic evaluation of known allosteric-site recovery in *apo* and substrate-bound structures across a broader range of proteins.

### AllositeUltra decouples site detection from preformed pocket geometry

These findings suggest that cross-state recognition requires moving beyond the assessment of currently visible pockets to the evaluation and discovery of potential allosteric regions across conformations. To this end, we developed AllositeUltra, which integrates information across scales through multimodal pocket scoring. Residue-guided candidate supplementation further extends the search to regions missed by initial geometric detection (Fig. 1g).

Existing methods often use geometric detection to define candidate pockets before evaluating their allosteric potential. Conformational changes may cause regions to be missed when they no longer meet geometric detection criteria, or alter detected pockets’ local features and affect subsequent scoring. Despite these limitations, AllositePro performed comparatively well across states in the preceding evaluation. AllositeUltra therefore retains its pocket-level representation framework while incorporating residue environments and protein-wide context to complement local recognition cues that may weaken with conformational change. The model combines SaProt’s structure-aware embeddings^28^ with residue physicochemical properties and dynamic responses to characterise the sequence–structure environment of candidate regions. It also uses SaProt’s inter-residue attention to represent associations between candidate regions and the rest of the protein. The outputs of the three branches are integrated into a site score (Supplementary Fig. 2).

Building on multimodal site evaluation, AllositeUltra uses residue-guided candidate supplementation to construct candidate regions from learned allosteric cues. Within the deep-learning branch, pocket classification, ranking and residue-level probability prediction are jointly optimised, allowing site evaluation and residue localisation to share representations. The model selects residues according to their predicted probabilities, surface accessibility and neighbourhood relationships, then spatially clusters them into supplementary candidates. These candidates are scored and ranked alongside the initial geometry-derived pockets, bringing residue-guided candidate generation back into the same site-evaluation framework. This mechanism allows residue representations to shape the candidate search space directly, extending site discovery beyond preformed cavities to potential allosteric regions indicated by residue predictions. It thus provides an alternative route to identifying cryptic sites that lack distinct pocket geometry in the input conformation (Supplementary Fig. 2).

### AllositeUltra achieves superior cross-state recognition

To assess whether AllositeUltra addressed the cross-state recovery limitations identified above, we evaluated it on AsBench2, with particular emphasis on *apo* and substrate-bound structures. We compared AllositeUltra with AllositePro, PASSER, PASSER2, PASSERRANK and DeepAllo^13,14,24,29–31^ across the allosteric ligand-bound, *apo* and substrate-bound test sets.

We assessed localisation accuracy relative to experimental allosteric sites using pocket-centre distance, minimum atom distance and key-residue coverage across Top-n cut-offs. AllositeUltra achieved shorter distances and higher coverage at most cut-offs across all three states, indicating improved site localisation and residue recovery across input conformations (Fig. 2a). We then applied a joint criterion requiring both spatial localisation and residue coverage to assess whether these improvements translated into successful site identification. AllositeUltra showed higher Top-n success rates overall than the comparison methods (Fig. 2a). Because experimental validation typically prioritises a small number of candidates, we focused on Top-3 success. AllositeUltra achieved 85.7% on the Core-set, exceeding the strongest baseline by 11.4 percentage points. AllositeUltra showed larger gains over the corresponding strongest baselines in the *apo* and substrate-bound states commonly available during discovery. Top-1 success was also highest across all three structural states (Fig. 2b). These results demonstrate improved site recovery across structural states, supporting the identification of potential regulatory sites from available conformations before an allosteric ligand-bound structure is known.

**Fig 2.**
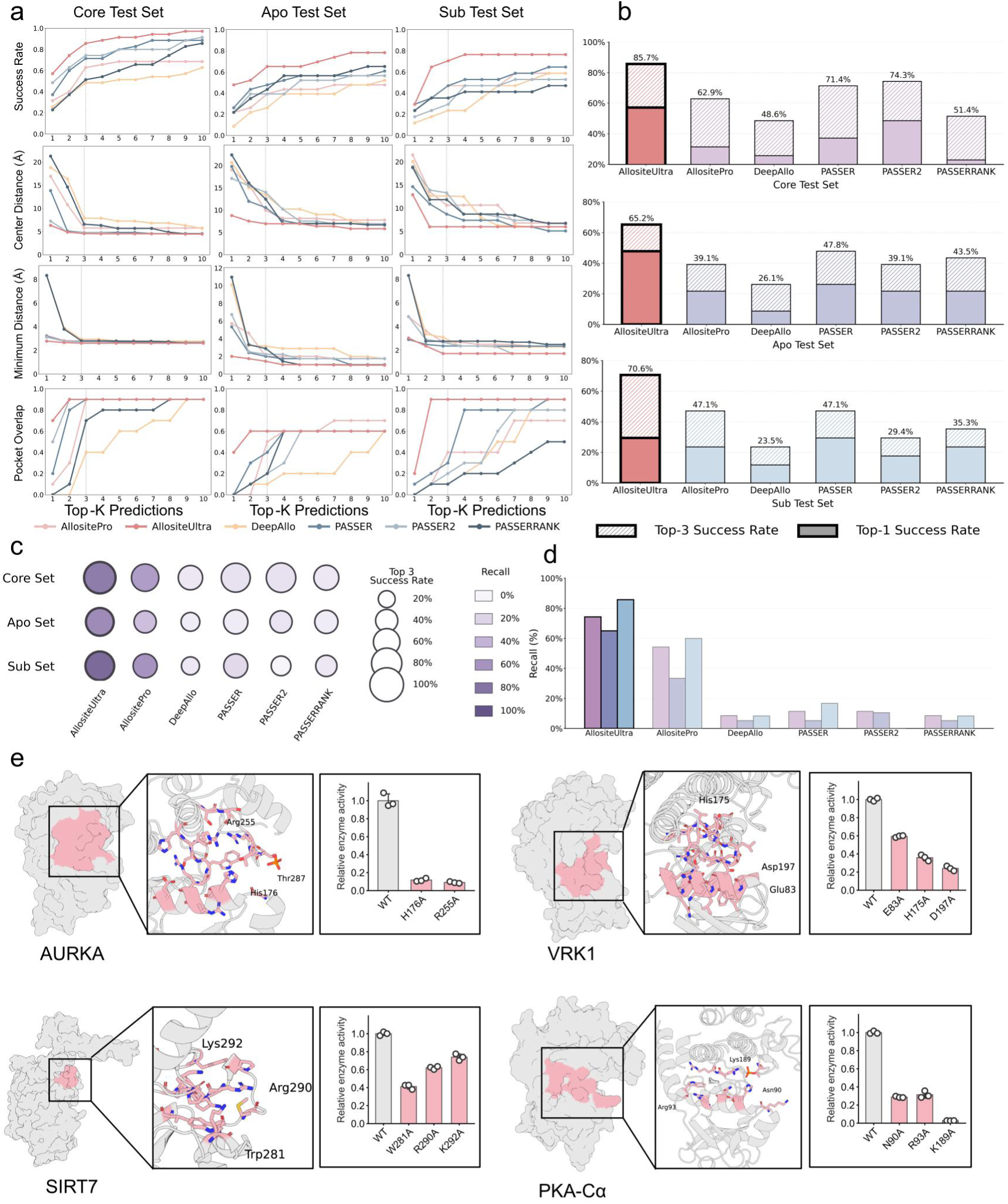
Cross-state prediction performance and functional validation of AllositeUltra-predicted pockets by mutagenesis. a, Top-n success rates, pocket-centre distances, minimum atom distances and reference-residue coverage of AllositeUltra and baseline methods on the allosteric ligand-bound, *apo* and substrate-bound test sets. The latter three metrics are shown as medians. b, Top-1 and Top-3 success rates, represented by solid segments and total bar heights, respectively. c, Joint comparison of Top-3 success and recall, represented by circle size and colour, respectively. d, Candidate-pocket recall. e, Predicted pocket locations and enzyme activities following pocket-residue substitutions in AURKA, VRK1, SIRT7 and PKA-Cα. Pockets are highlighted in pink, and activities are normalized to the corresponding wild-type control. Data are mean ± s.d.; each point represents one technical replicate well from a single experiment (n = 3 per condition).

Success among top-ranked candidates does not fully capture the risk of missed detections during prospective screening. Proteins without any predicted positive pockets may be excluded from subsequent analysis. We therefore measured candidate-level recall, defined as the proportion of site-matching candidates classified as positive. AllositeUltra achieved higher recall than all baselines across the three test sets (Fig. 2d). Together, Top-3 success and recall indicated improved prioritisation of experimental sites and retention of site-matching candidates as positive predictions across structural states (Fig. 2c).

Despite these improvements, site recovery on the Apo-set and Sub-set remained lower than in the allosteric ligand-bound state. To examine whether this performance gap was associated with local conformational changes, we grouped the paired samples included in this analysis by pocket RMSD. Although AllositeUltra achieved lower Top-3 success for large than small conformational changes, its rate in the large-change group was four times that of the strongest baseline. To examine the source of this advantage, we compared predictions before and after candidate supplementation. With only the initial candidates, the model already outperformed the strongest baseline in the large-change group. Residue-guided candidate supplementation doubled the success rate, indicating that supplementation helped alleviate recognition difficulties associated with larger local conformational changes. Refinement ablations showed consistent improvements across all three structural states. Integrating all three representation branches also yielded higher overall Top-3 success than single- or two-branch combinations, supporting their complementary contributions (Supplementary Fig. 3).

### Mutagenesis confirms the functional relevance of predicted pockets

Applying cross-state recognition to the discovery of new allosteric sites also requires testing the functional relevance of predicted pockets. To this end, we selected four previously unreported candidate allosteric pockets predicted by AllositeUltra in AURKA, PKA-Cα, VRK1 and SIRT7 and measured the effects of pocket-residue mutations on protein activity (Fig. 2e).

We first examined the extensively studied kinases AURKA and PKA-Cα. AURKA regulates mitosis through its roles in centrosome function, spindle assembly and chromosome alignment^32^. AllositeUltra predicted a candidate pocket at the junction of its αC helix, DFG motif and activation segment. Mutating His176 or Arg255 reduced substrate phosphorylation. PKA-Cα, the catalytic subunit of cAMP-dependent protein kinase, transmits cAMP signals through substrate phosphorylation^33^. The model identified another candidate pocket near its αC helix and neighbouring regulatory structures, where mutations of Asn90, Arg93 or Lys189 likewise reduced kinase activity. These results show that AllositeUltra can propose new functionally relevant candidate sites even in kinases with extensively characterized regulatory mechanisms (Fig. 2e).

We next examined VRK1, a nuclear serine/threonine kinase involved in chromatin regulation and the DNA damage response^34^. Its allosteric regulation is less extensively characterized than that of AURKA and PKA-Cα, making new activity-associated pockets useful starting points for investigating how VRK1 function can be modulated. AllositeUltra predicted a candidate pocket on the αC-helix surface and adjacent loops. Mutating Glu83, His175 or Asp197 reduced kinase activity, providing experimental evidence that this region contributes to VRK1 function (Fig. 2e).

These three systems tested kinases with different levels of prior regulatory characterization. To examine whether the functional relevance of predicted pockets extended to a different catalytic class, we selected the NAD⁺-dependent deacetylase SIRT7. SIRT7 contributes to chromatin and transcriptional regulation through deacetylation of histone and non-histone substrates^35^, using substrate-recognition and cofactor-dependent catalytic mechanisms distinct from those of protein kinases. The AllositeUltra-predicted pocket lies on the opposite face from the substrate-binding region, with the selected residues spatially separated from the NAD⁺-binding region. Mutating Trp281, Arg290 or Lys292 reduced deacetylase activity relative to wild type, supporting a functional link between the predicted pocket and SIRT7 catalysis (Fig. 2e).

### AllositeUltra enhances downstream ligand localization and pose prediction

To support allosteric drug discovery, site predictions should also guide downstream ligand localization and binding-pose modelling. To this end, we incorporated AllositeUltra-predicted pockets as spatial priors for ligand modelling and compared the resulting workflows with unguided modelling. The evaluation spanned conventional docking methods such as Vina and Glide^36,37^, AI-based pose prediction with DiffDock^38^, and co-folding approaches including Chai and Boltz^39,40^ (Fig. 3a).

**Fig 3.**
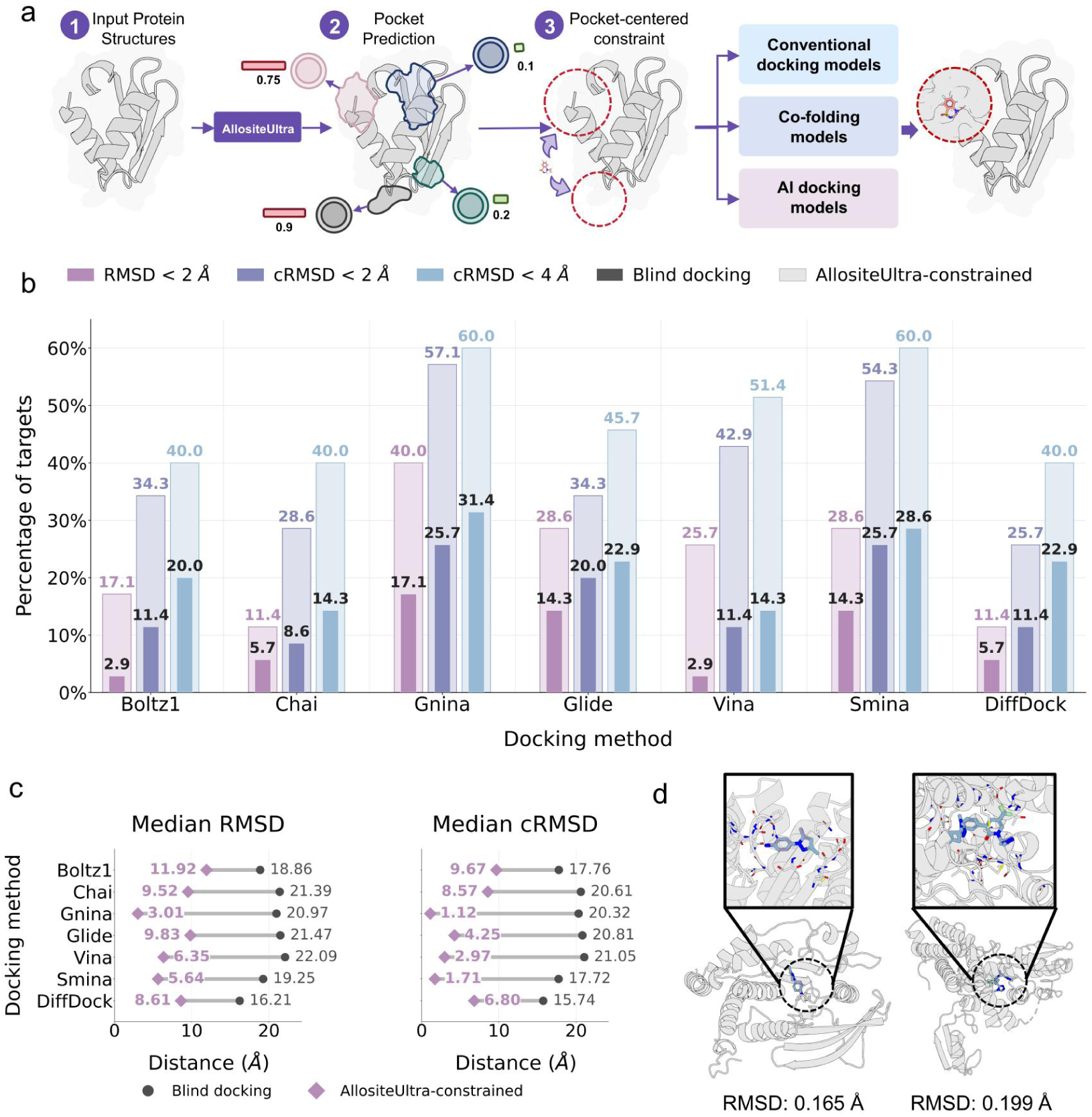
AllositeUltra improves allosteric ligand localization and binding-pose recovery. a, Workflow integrating AllositeUltra-predicted pockets into conventional docking, co-folding and AI-based docking models. b, Percentage of systems whose top-ranked pose achieves ligand heavy-atom RMSD < 2 Å or ligand-centre distance (cRMSD) < 2 Å or < 4 Å under unguided and AllositeUltra-guided conditions. RMSD accounts for ligand symmetry; centre distance measures the separation between predicted and experimental ligand heavy-atom centroids. Dark and light shades denote unguided and AllositeUltra-guided modelling, respectively. c, Median ligand RMSD and centre distance for the corresponding poses, with lines connecting results for the same method under both conditions. d, Representative superpositions of ligand poses predicted by Gnina with AllositeUltra guidance and their experimental conformations. From left to right: PTP1B (PDB: 8G65, chain A) and CBL-b (PDB: 8QTK, chain A), with ligand heavy-atom RMSDs of 0.165 and 0.199 Å, respectively.

AllositeUltra-guided workflows placed predicted ligands closer to the experimentally defined allosteric binding regions. For top-ranked predictions, AllositeUltra guidance reduced the median ligand-centre distance by 45.5–94.5% across the seven methods (Fig. 3c). The fraction of systems with a localization error below 4 Å increased across all methods, by 17.1–37.1 percentage points (Fig. 3b). Site guidance also improved binding-pose recovery, increasing the success rate at heavy-atom RMSD below 2 Å to 2.00–9.00 times that of unguided modelling (Fig. 3b) and yielding ligand poses nearly superimposable on experimental conformations in several representative systems (Fig. 3d). Top-three and top-five predictions showed consistent trends (Supplementary Fig. 4).

Having established improvements over unguided modelling, we assessed how closely predicted-site guidance approached guidance from experimentally defined sites. This comparison also included methods requiring a predefined binding region. With AllositeUltra guidance, nearly half of the methods achieved median centre distances below 4 Å, and approximately half were within 2 Å of the experimental-site reference. For pose recovery (RMSD < 2 Å), 70% of methods achieved at least 75% of the reference success rate (Supplementary Fig. 5). These results show that AllositeUltra can translate allosteric-site predictions into spatial guidance for ligand modelling, improving binding-region localization and pose recovery and providing a practical starting point for structure-guided allosteric ligand discovery.

### Prospective screening in MAGI2 uncovers a novel crystallographically validated site

The preceding evaluation showed that predicted pockets could guide the localisation and binding-pose modelling of known ligands. We next conducted a prospective study in MAGI2, applying the site-guided strategy to discover new ligands. MAGI2 is a membrane-associated scaffold containing multiple PDZ and WW domains and a guanylate kinase-like (GK) region. It helps organise receptor and adhesion-protein complexes at synaptic junctions. Its N-terminal PDZ0 and GK domains form a supramodule that recognises phosphorylated binding partners, including SAPAP proteins^41^. We focused on this supramodule to identify potential small-molecule binding sites outside the phosphopeptide-recognition groove.

AllositeUltra identified a candidate pocket at the PDZ0–GK interface of murine MAGI2, spatially separated from the phosphopeptide-recognition groove (Fig. 4b). The pocket lies between the PDZ0 β-sheet surface and a GK helix and comprises residues from both domains. Guided by the predicted site, we virtually screened a selected fragment library and prioritised 40 compounds for microscale thermophoresis (MST). Five compounds showed detectable binding (Supplementary Fig. 6 and Supplementary Table 3), including the micromolar-affinity initial hit MDL-6-H6, which had a dissociation constant of 25.9 μM (Fig. 4a).

**Fig 4.**
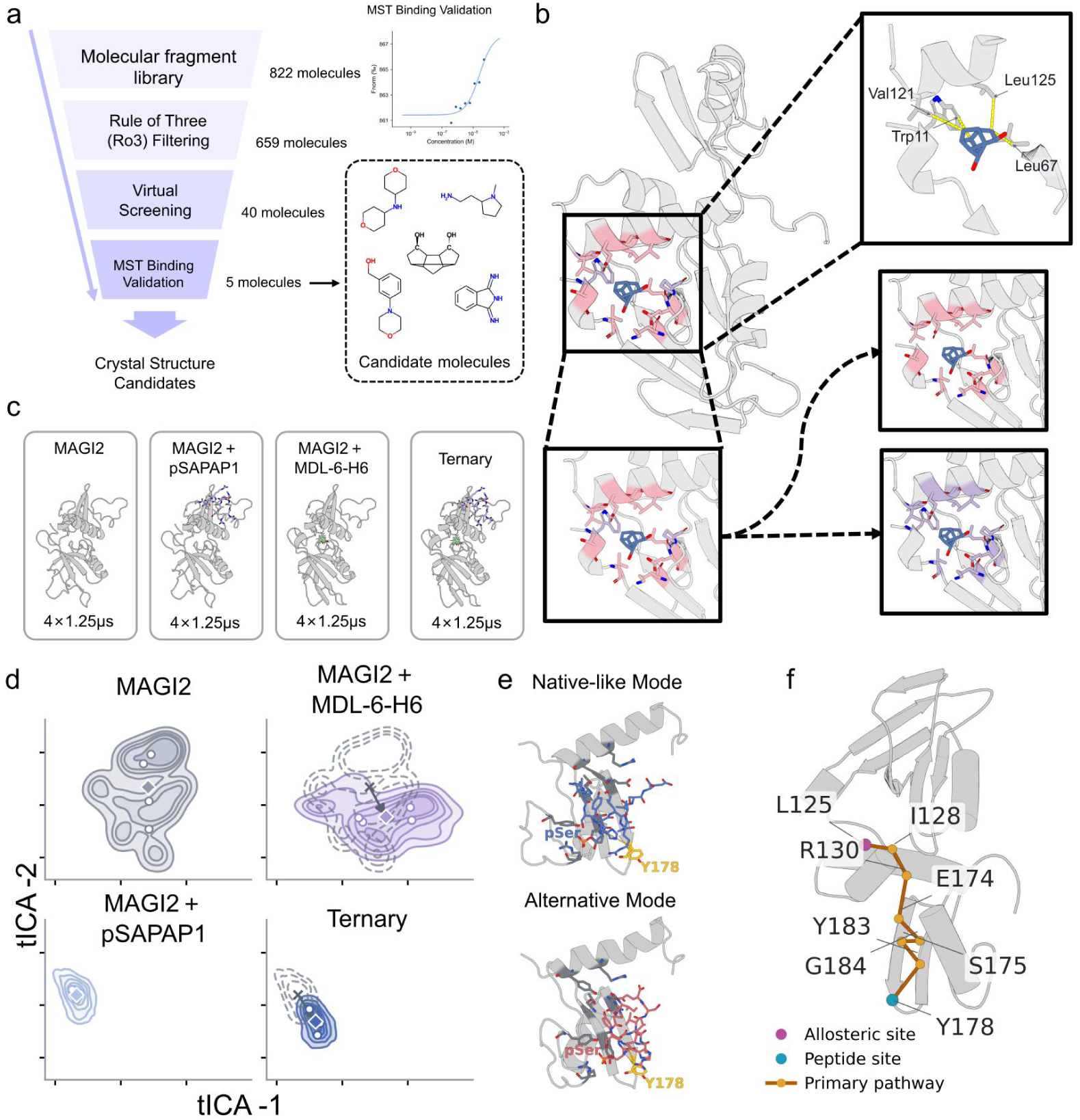
AllositeUltra-guided discovery and molecular dynamics analysis of a candidate allosteric site in MAGI2. a, Predicted-site-guided fragment screening and experimental validation, with candidate chemical structures and the microscale thermophoresis (MST) binding curve for MDL-6-H6. b, Crystal structure of the MAGI2 PDZ0–GK supramodule bound to MDL-6-H6 at 2.24 Å resolution (PDB: 26PA). Insets show the ligand-contact environment and correspondence between the predicted region and the crystallographic binding site. c, Four molecular dynamics systems: *apo* MAGI2, MAGI2–pSAPAP1, MAGI2–MDL-6-H6 and the ternary complex, each simulated in four independent 1.25-μs trajectories. d, Comparison of conformational distributions across the four simulated systems, showing shifts upon MDL-6-H6 binding through time-lagged independent component analysis (tICA). e, Representative phosphopeptide-binding modes: a native-like mode retaining the principal contacts of the experimental structure ^41^ and an alternative mode with a reorganized contact network. The relative positions of phosphoserine (pSer) and Tyr178 are highlighted. f, A potential allosteric pathway identified by residue-network analysis, linking Leu125 in the predicted pocket to the phosphopeptide-recognition region near Tyr178.

Crystallographic screening of all five binders yielded a 2.24 Å high-resolution crystal structure of MDL-6-H6 bound to the murine MAGI2 PDZ0–GK supramodule (PDB: 26PA; Supplementary Table 4). The structure defined the ligand’s binding position and conformation at the atomic level. Its carbon scaffold occupied a hydrophobic depression formed by Leu67, Val121 and Leu125 and contacted the aromatic side chain of Trp11 (Fig. 4b). The predicted pocket closely matched the experimental binding region, with pocket and ligand centres 1.50 Å apart and 80% coverage of ligand-contacting residues. Structural comparison showed that one Lys96 side-chain conformation in the phosphopeptide-bound state extended into the space occupied by MDL-6-H6. This occlusion may make a distinct cavity less apparent in the input structure and could lead geometry-based methods to miss the site or underestimate its binding potential. AllositeUltra nevertheless accurately localised the site in this conformation and guided new ligand discovery, demonstrating its cross-state recognition capability when local cavity cues were limited.

When evaluated against the experimental binding mode, site-guided docking recovered 90% of ligand-contacting residues. We performed molecular dynamics simulations to assess the persistence of key contacts during conformational fluctuations. Leu67, Val121 and Leu125, assigned high allosteric-site residue probabilities by AllositeUltra, each maintained ligand-contact occupancies above 94% and showed favourable contributions in residue-level energy decomposition. These simulations complemented the crystal structure by characterising the persistence of key contacts (Supplementary Fig. 7). Starting from a phosphopeptide-bound structure, this prospective study identified an initial hit with micromolar affinity. The 2.24 Å high-resolution complex crystal structure (PDB: 26PA) directly confirmed the new binding site identified by AllositeUltra and the ligand’s binding mode at the atomic level.

Following crystallographic localization of MDL-6-H6, we used molecular dynamics simulations to compare *apo* MAGI2, MAGI2–pSAPAP1, MAGI2–MDL-6-H6 and the ternary complex (Fig. 4c). All systems retained their overall folds, and MDL-6-H6 remained in the interface pocket (Supplementary Fig. 8). Compared with MAGI2–pSAPAP1, the ternary complex showed increased state 3 occupancy and decreased state 6 occupancy (Fig. 4d and Supplementary Fig. 9a). These shifts were accompanied by interface-contact rearrangements and an increased interdomain hinge angle (Supplementary Fig. 9). Contact-based clustering identified two phosphopeptide-binding modes: a native-like mode retaining the principal contacts of the experimental complex (PDB: 7YKI)^41^ and an alternative mode with reorganized contacts (Fig. 4e). In the ternary complex, the native-like population was 89.2 percentage points lower than in MAGI2–pSAPAP1. This shift coincided with reduced retention of crystallographic protein–peptide contacts and a shorter phosphoserine–Tyr178 distance. The phosphopeptide remained bound to MAGI2 while its interactions with the recognition region were reorganized. To complement this contact analysis, we compared phosphopeptide MM/GBSA endpoint interaction scores using two generalized Born (GB) models. Both models yielded less favourable scores in the ternary complex than in MAGI2–pSAPAP1 (Supplementary Fig. 10). We further combined inter-residue motion correlations with residue contacts in network analysis to examine dynamic links between the interface pocket and the distal phosphopeptide-recognition region. The network identified a potential allosteric pathway from Leu125 in the AllositeUltra-predicted pocket through the GK domain to the recognition region near Tyr178 (Fig. 4f). These results suggest that MDL-6-H6 binding allosterically alters phosphopeptide recognition by MAGI2.

### AllositeUltra maps the candidate allosteric landscape across the human proteome

In MAGI2, the predicted pocket guided new ligand discovery and was validated by crystallography. To extend site exploration from a single protein to a broader range of targets, we applied AllositeUltra to the human proteome. Integrating AlphaFold DB predictions^42^ with experimental PDB structures, we identified 26,366 candidate allosteric pockets across 12,252 human proteins (Fig. 5a). Only 26.2% of these proteins had existing drug-related records, indicating broad coverage of proteins lacking such annotations. At the site level, comparison with AllositePro^13^ identified 9,214 candidate pockets in shared regions and 13,407 outside AllositePro-predicted regions (Fig. 5b). The atlas thus broadened the scope of allosteric-site exploration at both the protein and pocket levels.

**Fig 5.**
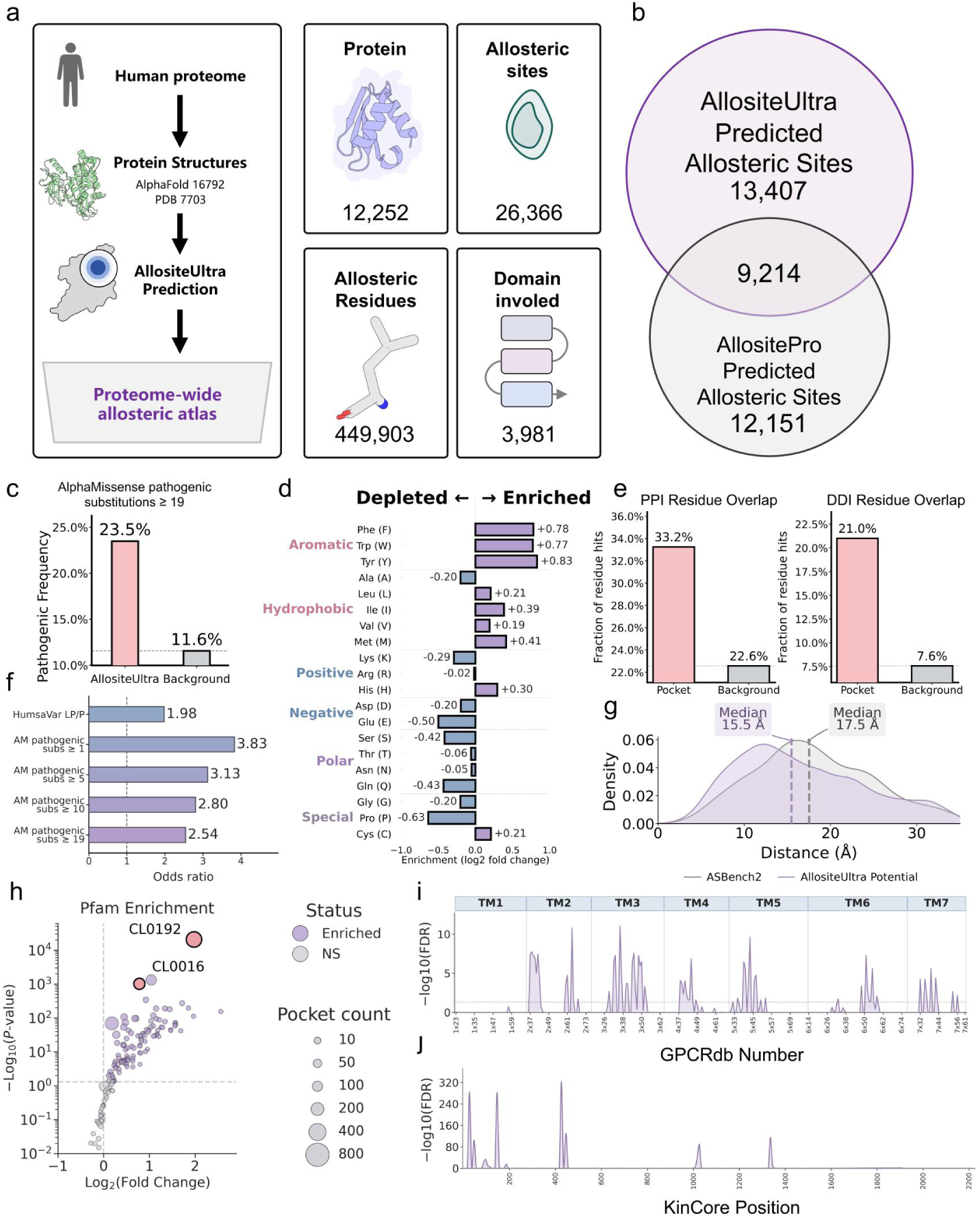
Mapping and characterization of candidate allosteric sites across the human proteome. a, Atlas construction workflow and coverage. b, Comparison of AllositeUltra- and AllositePro-predicted pockets in proteins evaluated by both methods. c, Frequency of disease-associated variants in candidate pockets and the corresponding protein-sequence background. d, Amino acid preferences of candidate pockets, expressed as log₂ fold changes relative to background. e, Overlap of candidate-pocket and background residues with protein–protein and interdomain interfaces. f, Enrichment of disease-associated variants based on Humsavar and AlphaMissense annotations ^43,46^, shown as odds ratios with 95% confidence intervals. g, Distributions of distances from candidate pockets and experimentally annotated AsBench2 sites to active-site residues; dashed lines indicate medians. h, Enrichment of candidate-pocket residues across Pfam clans. Circle size denotes pocket count, and colour distinguishes enriched from non-significant clans. Enrichment is defined by an unadjusted P < 0.05 from a one-sided hypergeometric test and fold enrichment > 1. i, Positional enrichment of candidate sites mapped to GPCRdb generic numbering ^44^, with transmembrane helices TM1–TM7 indicated above. j, Positional enrichment of candidate sites mapped to KinCore structural alignment coordinates ^45^. Both panels show enrichment significance as −log₁₀(FDR) after Benjamini–Hochberg correction.s

To characterize these regions and their potential regulatory context, we examined residue composition, structural location and disease-variant distribution. Candidate pockets were enriched in aromatic and several hydrophobic residues relative to the complete sequences of the corresponding proteins (Fig. 5d). Candidate-pocket residues overlapped protein–protein interaction and interdomain interfaces at frequencies of 33.2% and 21.0%, respectively, exceeding the corresponding backgrounds. This enrichment suggested that candidate regions tended to occur at intermolecular or interdomain contact sites (Fig. 5e). Their median distance to known functional centres was 15.5 Å, with a distribution similar to experimentally supported allosteric sites in AsBench2 (Fig. 5g). Disease-associated variants were enriched at candidate-pocket residues relative to the complete sequences of the corresponding proteins. This enrichment was supported by both known variant annotations from Humsavar and computational pathogenicity predictions from AlphaMissense^43^, and remained consistent across different AlphaMissense filtering thresholds (Fig. 5c,f).

We analysed the distribution of candidate pockets across Pfam clans^27^, identifying clans enriched in candidate-pocket residues to characterize their overall structural context (Fig. 5h). To resolve candidate-site positions within individual families, we focused on GPCRs and protein kinases, two major protein families. We mapped candidate sites onto their respective generic structural coordinates. GPCRdb mapping^44^ revealed recurrent candidate pockets around TM2, TM3 and TM5, whereas KinCore analysis^45^ showed preferential localization near the catalytic loop and DFG and HRD motifs (Fig. 5i,j). Clan-level enrichment and family-level positional mapping characterized the structural contexts and recurrent regions of candidate sites, providing leads for subsequent ligand screening and functional validation.

## Discussion

Our results showed that allosteric-site recovery varied with the input conformation, limiting how well ligand-bound benchmark performance reflected site recovery under prospective discovery conditions. AsBench2 linked experimentally supported sites to *apo* and substrate-bound structures, providing a basis for systematically examining this dependence. AllositeUltra combined multimodal pocket scoring with residue-guided candidate supplementation, improving recovery from conformations available during discovery.

Cross-state recognition raises a broader scientific question for structure-based prediction: how can available conformations reveal regions that support molecular binding in other states? Conformational changes alter local geometry, surface exposure and interaction environments^9^, limiting how fully a pocket or interface in the available structure reflects its binding potential. This question warrants investigation in predictions of other ligand-binding sites and protein–protein interaction interfaces. Such studies could test whether models can locate candidate sites before their binding geometry is fully exposed, supporting molecular discovery when bound structures are unavailable.

The prospective MAGI2 study illustrates a practical route from cross-state recognition to molecular discovery. A pocket predicted from the phosphopeptide-bound structure guided screening of a fragment library, yielding MDL-6-H6 as an initial hit with a dissociation constant of 25.9 μM. The high-resolution crystal structure of the complex, determined at 2.24 Å (PDB: 26PA), revealed the atomic details of MDL-6-H6’s binding mode in the interface pocket. The observed binding position closely matched the prediction, providing direct crystallographic confirmation that the model accurately localised this previously uncharacterised site. Building on this prospective validation, we applied AllositeUltra across the human proteome to systematically identify potential allosteric regions and construct an atlas of candidate sites. By integrating structural context, disease-associated variants and protein family information, the atlas informs candidate prioritisation across targets, supporting subsequent ligand screening and functional validation.

To further support ligand screening and design at candidate sites, future work could combine site recognition with protein conformational sampling. This would allow potential binding regions and conformations capable of accommodating ligands to be explored together. Within this framework, site predictions could provide spatial guidance for sampling candidate regions and associated structural elements. Local pocket optimisation could refine these conformations, providing structural models of potential binding environments. Candidate sites could then be rescored and ranked in these conformations to guide conformational selection, establishing feedback between site recognition and conformational exploration. Building on the current cross-state evaluation, such frameworks could be assessed against experimental ligand-bound structures for both site-localisation accuracy and recovery of local pocket conformations. The goal is to identify potential allosteric sites while providing protein conformations suitable for ligand modelling, connecting site discovery more directly to drug screening and design.

## Methods

### AsBench2 dataset construction and cross-conformation benchmark assembly

#### Construction of the AsBench1 paired challenge set

To evaluate cross-conformation allosteric-site prediction, we first selected AsBench1 Core-set entries with an identifiable pocket in the allosteric ligand-bound state. Entries were retained if Fpocket generated at least one candidate ^47,48^ with a pocket-centre distance ≤8 Å, a minimum atom distance ≤4 Å and a binding-site residue overlap ≥0.6, yielding 153 entries. We then focused on transferases, hydrolases and oxidoreductases, which offered larger sample sizes and broader structural-state coverage, and newly collected *apo* and substrate-bound experimental structures for the corresponding targets. These structures were paired with their allosteric ligand-bound counterparts in the Core-set, excluding duplicate structures. The resulting challenge set comprised 35 *holo*–*apo* pairs and 40 *holo*–substrate pairs, involving 59 distinct targets.

#### Data sources and experimental evidence for AsBench2

Starting from ASD2023 ^22^, we updated and expanded records of allosteric proteins and their modulators through literature searches and retrieved the corresponding complex structures from the PDB. Data collection was completed in October 2024. Original research articles were manually reviewed to verify experimental support for the functional effects of the modulators on their corresponding proteins. Relationships among proteins, modulators, allosteric sites and structural records were curated for subsequent quality filtering and dataset construction.

#### Core-set filtering and redundancy removal

AsBench2 curation followed the site-quality criteria of AsBench^23^. We included crystal structures containing non-covalent organic small-molecule modulators, excluding complexes with allosteric ions or peptides. Allosteric sites were required to be spatially distinct from substrate-binding sites, with no missing residues within 4 Å of the modulator. Redundant complexes were consolidated at the allosteric-site level, retaining one representative structure per site and treating distinct sites within the same protein separately. The resulting Core-set comprised 536 allosteric sites across 446 proteins.

#### Construction of the Apo-set and Sub-set

Using the Core-set as a reference, we collected corresponding experimental structures to construct the Apo-set and Sub-set. The Sub-set included structures bound to substrates, substrate analogues or other orthosteric ligands. In both sets, the target allosteric site was required to be unoccupied by a ligand. Correspondence with Core-set proteins was established through sequence and structural comparisons, considering construct consistency and structural completeness around the allosteric and substrate-binding regions. Structures were counted by unique PDB ID–chain combinations within each state, yielding 387 Apo-set structures and 275 Sub-set structures.

#### Training, validation and test splits

Before partitioning, we selected Core-set entries with a detectable cavity at the experimentally annotated allosteric site that satisfied the predefined geometric matching criteria. This filter was applied to the allosteric ligand-bound structure; corresponding *apo* and substrate-bound structures were included without requiring a detectable cavity at the mapped site. We partitioned entries by the date of the first experimental report of each allosteric site, using 1 January 2022 as the cutoff. Earlier sites were assigned to training and validation, and eligible sites first reported from the cutoff through October 2024 were considered for testing. All structures associated with a site were assigned to the same partition, irrespective of the release dates of their *apo* or substrate-bound structures. Core test chains were aligned with training chains using sequences extracted from structural coordinates. Pairs with ≥90% sequence identity and ≥90% coverage of each chain underwent site-level comparison after superposition of corresponding Cα atoms. Sites were considered overlapping when ligand heavy-atom centroids were within 8 Å and the Jaccard index of the corresponding 6 Å pocket-residue sets was ≥0.50. Overlapping Core test sites and their corresponding Apo/Sub entries were excluded, yielding 35 Core, 23 Apo and 17 Sub test entries (75 target–state records).

#### Construction of the AsBench-AFDB pretraining set

To expand the pretraining data, AsBench2 Core-set structures were queried against the human proteome in AlphaFold DB using Foldseek. Candidates with a matching probability ≥0.8 were aligned using TM-align, and known allosteric sites were mapped onto the candidate structures. Mappings with severe spatial clashes were excluded. Mapped regions were required to contain at least five protein residues within 8 Å of the experimental ligand centre and at least five within 6 Å of the ligand atoms, and to overlap at least one Fpocket-predicted pocket. The retained structures and site annotations formed AsBench-AFDB for pretraining the deep-learning branch. Data corresponding to test-set sites were excluded from pretraining.

#### Training resampling

During fine-tuning, Core-set allosteric sites served as the basic sampling units, together with their corresponding Apo-set and Sub-set structures. Core-set entries were resampled at equal frequency to reduce training bias arising from uneven structural-state coverage, while retaining the available cross-state representations of each site.

#### Dataset composition and coverage analyses

The complete AsBench1 Core-set served as the reference for comparing dataset size and domain coverage with AsBench2. Entries were deduplicated by PDB ID and chain identifier within each dataset, and protein-chain sequences were annotated using PfamScan ^27^ to identify families and clans. A domain was considered to overlap an allosteric site if its envelope interval contained at least one protein residue within 6 Å of a modulator heavy atom. Structure-aware residue representations were extracted using SaProt-650M-PDB ^28^ and mean-pooled to obtain a 1,280-dimensional vector for each protein chain. Vectors from both datasets were combined for joint two-dimensional t-SNE embedding with a random seed of 42. Kernel density estimates were calculated from the resulting coordinates to visualize distributions in protein representation space.

### AllositeUltra model architecture

#### Overall workflow and initial candidate-pocket generation

AllositeUltra takes a protein structure as input and uses Fpocket4 ^47,48^ with default parameters to generate initial candidate pockets, recording their residue composition and extracting geometric and physicochemical descriptors. Candidates are scored by the deep-learning, Attention and machine-learning branches, whose outputs are combined into a pocket-level prediction score. Residue-level probabilities from the deep-learning branch are also used to generate supplementary candidate regions, which are merged with the initial pockets for joint scoring and ranking.

#### Deep-learning branch

The deep-learning branch uses SaProt-650M-PDB to extract a 1,280-dimensional structure-aware representation for each residue and integrates residue-level B-factors, Kyte–Doolittle hydrophobicity indices, solvent-accessible surface area (SASA) calculated using DSSP, and dynamic-response z-scores generated by Essential Site Scanning Analysis (ESSA) ^49^. SaProt representations are projected to 256 dimensions through a linear layer and concatenated with these features. A four-head self-attention module models inter-residue relationships to produce 64-dimensional representations. Pocket-level and residue-level decoders each use a four-head attention module followed by a multilayer perceptron to predict pocket allosteric probabilities and residue-level site probabilities. The branch jointly optimizes pocket classification, pocket ranking and residue classification through multitask learning.

#### Attention branch

The Attention branch extracts inter-residue attention matrices from all 33 SaProt layers, each containing 20 heads ^28^, after removing special tokens. For the attention matrix *A^l^*^,h^ from layer *l* and head h , the feature for candidate pocket *P* is the total attention directed from all protein residues to pocket residues:

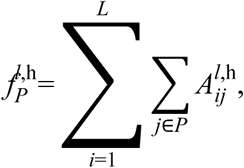

where *L* is the number of residues in the protein chain. These sums form a 660-dimensional pocket feature vector without normalization by pocket size or averaging across layers and heads. XGBoost learns nonlinear combinations of these features to predict pocket allosteric probabilities. Attention features are recalculated in the same way for refined pockets.

#### Machine-learning branch

The machine-learning branch retains the AllositePro descriptor framework ^13^, using the normalized maximum distance between α-spheres, hydrophobicity density and flexibility score as inputs to logistic regression:

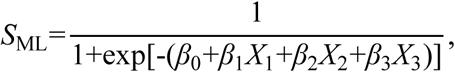

where *X*_1_ , *X*_2_ and *X*_3_ denote the three descriptors, *β*_0_ is the intercept, and *β*_1_ –*β*_3_ are the regression coefficients.

#### Integration of branch scores

The three branch outputs are combined by a weighted sum, with weights selected according to validation-set performance:

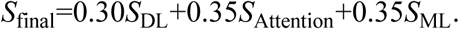

Candidate pockets are initially ranked in descending order of the composite score.

#### Residue-guided candidate-pocket refinement

Surface residues with predicted probabilities >0.5 are selected as candidates. Surface residues are defined as those with SASA >20 Å² or with at least three residues satisfying this SASA criterion within 8 Å. Surface residues with more than five high-confidence residues in their 6-Å Cα neighbourhood are also included. Selected residues are clustered by their three-dimensional coordinates using K-means, with 7– 35 residues per cluster. Clusters with centres separated by less than 8 Å are merged. Supplementary pockets comprise residues within 8 Å of each cluster centre; pockets containing fewer than 20 residues are expanded to 25 residues by adding the nearest residues.

Descriptors for supplementary pockets are extracted using Dpocket ^47^ (mode=E, d=8 Å) and Min–Max normalized against the Fpocket candidates from the same protein. Flexibility descriptors are rescaled using the normalization mapping derived from Fpocket outputs. Supplementary and initial candidates are pooled, features required by all three branches are calculated, and composite scores are obtained for the combined candidates. Cross-source redundancy is removed, prioritising candidates with higher initial fused scores, and candidate information is integrated to determine the final scores and ranking.

### Model training and ablation studies

#### Multitask loss function

The deep-learning branch jointly optimizes pocket classification, pocket ranking and residue classification. The total loss is defined as:

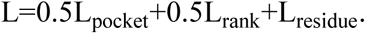

Pocket and residue classification use weighted binary cross-entropy, with positive-class weights of 20 and 10, respectively, and negative-class weights of 1. Pocket ranking uses a pairwise loss ^50^ between positive and negative candidates within the same protein:

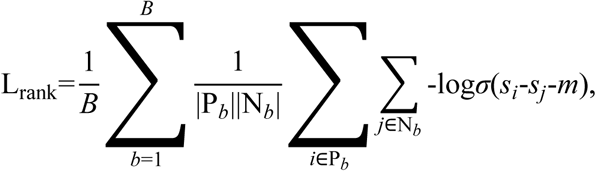

where *B* is the number of proteins in a batch, P*_b_* and N*_b_* are the positive and negative pocket sets, *s_i_* and *s_j_*are prediction scores, *σ* is the sigmoid function, and the ranking margin is *m*=0.3.

#### Training settings and pretraining strategy

The deep-learning branch was trained in two stages: pretraining and fine-tuning. SaProt parameters were frozen, and residue representations were extracted offline; only the downstream feature-processing layers, Transformer and prediction heads were trained. The model was implemented in PyTorch and optimized using Adam with a learning rate of 10^-4^, weight decay of 10^-5^ and batch size of 32.

AsBench-AFDB was divided into training and validation sets at an 80%/20% ratio using a random seed of 42. Pretraining ran for 100 epochs without early stopping, retaining the three highest-scoring checkpoints according to validation pocket Top-3 success rate after each epoch. The model was then fine-tuned on AsBench2 using the data partitioning and equal-frequency site resampling described above. Fine-tuning ran for up to 200 epochs, with model selection based on validation pocket Top-3 success rate and early stopping after ten consecutive epochs without improvement.

During pretraining, approximately 20% of valid residues were randomly selected in each protein, excluded from valid attention key/value positions, and assigned zero-valued fused features. This masking setting also applied during pretraining-stage validation and testing. Random masking was disabled during fine-tuning.

The Attention branch used SMOTE to balance positive and negative pockets at a 1:1 ratio, followed by an XGBoost classifier with binary logistic loss, ten trees, a maximum depth of five and a learning rate of 0.1. For the machine-learning branch, SMOTE with a random seed of 42 adjusted the minority-class count to 50% of the majority-class count before logistic regression was fitted.

#### Ablation study design

Component contributions were assessed through branch-combination and candidate-pocket refinement ablations. Branch ablations used the three branch scores produced by the trained model to construct single-and two-branch predictions without retraining. After a branch was removed, the remaining original weights were normalized to sum to one; single-branch configurations used a weight of one. Pocket rankings were recalculated for each configuration, and composite scores >0.5 were classified as positive predictions.

Refinement ablation compared predictions before refinement and candidate post-processing with the final predictions, while keeping the trained model unchanged. Both types of ablation were evaluated on the corresponding Core-set, Apo-set and Sub-set test subsets. Overall success rates were calculated across target–state records pooled from the three test subsets.

### Benchmark evaluation methods

#### Comparison methods and structure preprocessing

AllositePro, PASSER, PASSER2, PASSERRANK and DeepAllo were selected as comparison methods and run with the default parameters of their publicly available implementations. All methods received the same protein structures, processed using Gemmi to remove small-molecule ligands, solvent and non-standard amino-acid atoms. Protein-chain sequences required by DeepAllo were extracted from the processed structural coordinates. The AsBench1 paired challenge set was evaluated using DeepAllo, AllositePro and PASSERRANK, whereas the AsBench2 benchmark included all five comparison methods.

#### Geometric site matching and success criteria

Pocket localization was evaluated using pocket-centre distance, minimum atom distance and reference-residue coverage. Pocket-centre distance was defined as the Euclidean distance between the geometric centre of the predicted pocket residues, *C_P_*, and the geometric centre of the experimental allosteric modulator atoms, *C_L_*:

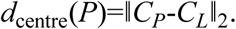

Minimum atom distance was defined as the shortest distance between a predicted pocket atom and a modulator atom:

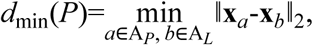

where A*_P_* and A*_L_* denote the pocket and modulator atom sets, respectively, and **x** denotes atomic coordinates. The ten protein residues closest to the allosteric modulator in the experimental structure formed the reference set K . Coverage was defined as:

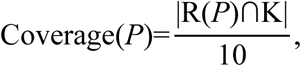

where R(*P*) is the predicted pocket residue set. A candidate was considered a successful match when the same pocket simultaneously satisfied *d*_centre_≤8 Å, *d*_min_≤4 Å and Coverage≥0.6. These criteria were applied consistently to candidate-detectability screening and cross-state evaluation.

#### Prediction performance metrics

Evaluation used the final candidate ranking returned by each method. A sample was counted as a Top-n hit if at least one of its first *n* candidates satisfied the joint success criteria. Top-n success rate was defined as:

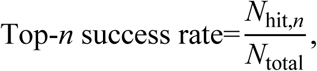

where *N*_hit,*n*_ is the number of samples with a hit among the first *n* candidates and *N*_total_ is the total number of evaluated samples. To assess localization accuracy across rank cut-offs, a single best pocket was selected from the first *n* candidates at each Top-n cut-off, and its centre distance, minimum atom distance and reference-residue coverage were calculated. Geometric metrics were summarized across samples using the median, 75th percentile and 90th percentile.

Recall was calculated from each method’s candidate-level classifications. Scores >0.5 defined positive predictions for AllositeUltra, AllositePro, PASSER, PASSER2 and DeepAllo, whereas PASSERRANK used scores >0. Recall was the fraction of candidates satisfying the joint site-matching criteria that were classified as positive:

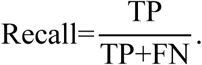

Here, TP and FN denote site-matching candidates classified as positive and negative, respectively, pooled across the evaluated samples.

#### Conformation-stratified evaluation

To assess the relationship between local conformational changes and cross-state recognition, we paired Core-set test sites with their corresponding *apo* or substrate-bound test structures. Duplicate pairs were removed, and different target states were counted separately. Reference pockets comprised residues in the designated Core chain whose minimum heavy-atom distance to the specified ligand was ≤6 Å. Residue correspondences were established using SIFTS; when unavailable, consensus mappings from all optimal global affine-gap sequence alignments were used. Global Kabsch superposition used matched Cα atoms across the full chains, and pocket Cα RMSD was calculated using the resulting transformation. Pairs required at least three non-collinear matched pocket Cα atoms and matched Cα coordinates for ≥80% of reference pocket residues. The 38 eligible pairs were pooled across target states and stratified using a fixed exploratory cutoff of 2 Å. This yielded a small-change group (<2 Å, 29 pairs) and a large-change group (≥2 Å, 9 pairs). We compared target-state Top-3 success rates across six methods within these groups. With the model and fusion scoring unchanged, we also compared the initial candidates with the complete candidate set after residue-guided supplementation but before post-processing. Group-wise success rates followed the Top-3 criteria defined above, using all pairs within each group as the denominator.

### Site-directed mutagenesis and functional assays

#### Protein expression and purification

Full-length human VRK1, PKA and SIRT7 coding sequences were cloned into pET28a, and human Aurora A (AURKA, residues 122–403) was cloned into pET30b. Site-directed mutagenesis generated VRK1 E83A, H175A and D197A; PKA-Cα N90A, R93A and K189A; SIRT7 W281A, R290A and K292A; and AURKA H176A and R255A. Wild-type and mutant plasmids were transformed into Escherichia coli BL21(DE3) for expression.

VRK1 was purified by nickel-affinity chromatography and dialysed into buffer containing 50 mM Tris-HCl (pH 8.0), 150 mM NaCl, 1 mM DTT and 5% glycerol. PKA was purified by nickel-affinity and size-exclusion chromatography and exchanged into storage buffer containing 50 mM Tris-HCl (pH 7.5), 150 mM NaCl, 0.25 mM DTT, 0.1 mM EGTA, 0.1 mM EDTA and 5% glycerol. SIRT7 was sequentially purified by nickel-affinity, cation-exchange and size-exclusion chromatography, then exchanged into storage buffer containing 150 mM NaCl, 50 mM K₂HPO₄ (pH 7.20), 1 mM DTT and 5% glycerol. AURKA was sequentially purified by nickel-affinity, anion-exchange and size-exclusion chromatography and exchanged into the same buffer used for VRK1.

#### SIRT7 activity assay

SIRT7 activity was measured using a Fluor de Lys (FDL) assay in Costar 3915 96-well plates (Corning, USA). Reactions were performed in a total volume of 75 μL in buffer containing 50 mM HEPES-Na (pH 7.4), 100 mM KCl, 200 μM TCEP and 1 mg/mL BSA. Reactions contained 100 μM Ac-APRK(dec)-AMC substrate, 12.5 mM NAD⁺ and 500 nM wild-type or mutant SIRT7 and were incubated at 37°C for 2 h. Stop solution containing 5 mg/mL trypsin and 4 mM nicotinamide was then added, followed by incubation at room temperature for 1.5 h to release the AMC fluorophore.

#### Kinase activity assays

PKA-Cα, AURKA and VRK1 activities were measured using the ADP-Glo Kinase Assay kit in opaque white 96-well plates (FCP968, Beyotime, China). Reactions were performed in a total volume of 30 μL. PKA-Cα and AURKA assays used buffer containing 40 mM HEPES (pH 7.5), 20 mM MgCl₂, 0.1 mg/mL BSA and 50 μM DTT. VRK1 assays used buffer containing 40 mM Tris-HCl (pH 7.5), 20 mM MgCl₂, 0.1 mg/mL BSA, 50 μM DTT and 2.5 mM MnCl₂.

For PKA-Cα, 6 μL of compound working solution was pre-incubated with 12 μL of enzyme solution at room temperature for 30 min. DMSO was introduced via the substrate peptide, resulting in a final concentration of 5% (v/v) in the reaction. Reactions were initiated by adding 12 μL of substrate mixture and incubated at room temperature for 40 min, with final concentrations of 7.5 nM enzyme, 100 μM kemptide and 25 μM ATP. AURKA reactions contained 40 nM enzyme, 100 μM kemptide and 25 μM ATP, whereas VRK1 reactions contained 750 nM enzyme, 100 μM H3tide and 50 μM ATP. Both AURKA and VRK1 reactions contained a final DMSO concentration of 5% (v/v), introduced via their respective substrate peptides, and were incubated at 37°C for 1 h.

ADP-Glo reaction termination and luminescence detection were performed according to the manufacturer’s instructions. Luminescence was measured using a Revvity microplate reader and normalized to the corresponding wild-type control.

Substrate peptides were synthesized by GL Biochem (China) Ltd.: Ac-APRK(dec)-AMC for SIRT7, kemptide (LRRASLG) for PKA-Cα and AURKA, and H3tide (ARTKQTARKSTGGK) for VRK1.

For the SIRT7, PKA-Cα, AURKA and VRK1 activity assays, each condition was measured in three technical replicate wells within a single experiment, and signals were normalized to the corresponding wild-type control.

### Site-guided ligand modelling

#### Evaluation systems and input preparation

Evaluation used 35 non-redundant protein–allosteric ligand systems from the test set described above. Receptors were obtained by removing ligands from experimental complexes while retaining the ligand-bound protein coordinates. Ligands were extracted from the corresponding experimental complexes, converted to canonical SMILES after bond-order correction, and prepared in the formats required by each modelling method. Experimental complexes were used to define experimental-site references and evaluate predicted poses.

#### Modelling methods and site-guidance settings

Vina, Smina ^51^, Gnina ^52^, Glide, DiffDock, Chai-1 and Boltz-1 were compared under three conditions: unguided modelling, AllositeUltra-predicted-site guidance and experimental-site guidance. Unguided modelling received no site information. AllositeUltra guidance used the three highest-ranked predicted sites, with separate modelling runs for each site followed by pooling of candidate poses. For these seven methods, predicted site centres were calculated by averaging representative residue coordinates, using Cα for glycine and the side-chain heavy-atom centroid for other residues. Experimental-site guidance used the experimental ligand heavy-atom centroid to define the search region or select restraint residues.

For Vina, Smina and Gnina, unguided search regions encompassed the entire receptor with a 4-Å extension in each direction. Site-guided runs used 20 × 20 × 20 Å³ boxes centred on the corresponding sites, with exhaustiveness set to 32. Smina used the Vina scoring function, whereas Gnina v1.3.2 used its default convolutional neural network to rescore poses after Vina-based conformational search. Glide 9.1 used SP mode and the OPLS_2005 force field. Unguided runs used cubic grids centred on the receptor heavy-atom centroid, with outer and inner box dimensions of 134 Å and 67 Å, respectively; site-guided runs used outer and inner dimensions of 20 Å and 10 Å. These four methods retained up to 40 poses per search region. Unguided DiffDock runs used DiffDock-L v1.1 ^38,53^, whereas site-guided runs used DiffDock-Pocket v1.0.0 ^54^ with its native pocket-centre interface and pocket cropping with a 10-Å buffer. Native flexible-side-chain treatment was retained without hard pose restraints, and five candidates were generated per run. Chai-1 (chai_lab 0.3.1) and Boltz-1 (boltz 2.2.0, model set to boltz1) used their native pocket-restraint interfaces, selecting the three mappable receptor residues nearest each site centre. Chai-1 applied soft ligand–residue restraints of 0–8 Å through its native POCKET interface. Boltz-1 designated the ligand as the binder in its native pocket constraint, set the maximum distance to 6 Å and enabled forced constraints. Both methods generated five candidates per site.

Experimental-site controls followed the corresponding site-guided workflows, replacing predicted site centres with experimental ligand centres. Under AllositeUltra guidance, candidates sampled separately at the three predicted sites were pooled for joint scoring and ranking. The candidate limits specified above applied to each search region or site, and no additional poses were added when fewer candidates were returned.

PocketXMol ^55^, QuickVina2^56^ and SurfDock ^57^ were compared under predicted-site and experimental-site guidance, using the three highest-ranked AllositeUltra sites for the predicted condition. PocketXMol used the official Gaussian/free configuration without flexible-receptor treatment, a 15-Å pocket radius supplied through pocket_coord, and ETKDGv3^58^ conformers translated to each site centre. Experimental-site runs used the experimental ligand heavy-atom centroid. Sampling used 100 diffusion steps and 100 poses per site and seed, ranked by the official tuned confidence model.

QuickVina2 used 20 × 20 × 20 Å³ search boxes and thread=8000. Predicted centres were calculated from all available pocket-residue heavy atoms; experimental centres were obtained from the complex ligand. Up to 20 poses per site were imported and ranked by Vina affinity. SurfDock constructed receptor surfaces within 8 Å of independently generated conformers translated to predicted centres, with the site centre supplied to the model; experimental-site runs used experimental ligand coordinates for surface and pocket construction. SurfDock used Screen mode, 20 steps and 40 poses per site and seed, ranked by pose confidence. PocketXMol and SurfDock used seeds 42, 43 and 44.

#### Evaluation of ligand localization and binding poses

Candidate poses were ranked using each method’s native score. Candidates from the three AllositeUltra-predicted sites were pooled and ranked jointly, with the Rank-1 pose used for the primary evaluation. Supplementary analyses considered the three and five highest-ranked candidates, taking the minimum ligand-centre distance and minimum heavy-atom RMSD separately. All success rates were calculated using all 35 systems as the denominator. Systems without evaluable poses or valid values for the corresponding metric were counted as failures for that criterion. Medians were calculated across systems with valid values for each metric, with the corresponding number of evaluable systems reported separately for each metric, method, guidance condition and Top-k cut-off. Before evaluation, corresponding Cα atoms in the predicted structure and experimental complex were superposed using the Kabsch algorithm, and the same rigid-body transformation was applied to the predicted ligand. Centre distance was defined as the Euclidean distance between predicted and experimental ligand heavy-atom centroids. Ligand heavy-atom RMSD was calculated using spyrmsd 0.9.0 ^59^, with molecular graph isomorphism accounting for atom correspondence and ligand symmetry. No independent ligand superposition was performed after receptor alignment.

### Virtual screening and experimental validation of MAGI2 ligands

#### Compound library preparation and virtual screening

The initial library of 822 compounds was provided by the National Center for Protein Science (Shanghai). Compounds were filtered using an extended Rule of Three (Ro3) ^60^, retaining those with a molecular weight ≤300 Da, no more than three hydrogen-bond donors and three acceptors, cLogP or cLogD ≤3, no more than three rotatable bonds, and a topological polar surface area ≤60 Å². The resulting library contained 659 compounds. Virtual screening was performed using Glide, targeting the MAGI2 PDZ0–GK interface pocket predicted by AllositeUltra. Docking scores, MM/GBSA binding-energy estimates and predicted binding modes were jointly considered to select 40 candidate compounds for subsequent microscale thermophoresis binding measurements.

#### MAGI2 PDZ0-GK Protein Purification

The gene fragment encoding the murine MAGI2 PDZ0-GK superdomain was cloned into the pET.M.3C prokaryotic expression vector. The recombinant plasmid was transformed into Escherichia coli BL21 (DE3) competent cells. A single colony was inoculated and grown in LB medium at 37 ℃ with shaking until the OD600 reached 0.8–1.0. Protein expression was induced by adding 0.2 mM isopropyl β-D-1-thiogalactopyranoside (IPTG), followed by incubation at 16 ℃overnight with shaking. Cells were harvested by centrifugation and resuspended in lysis buffer (25 mM Tris, pH 8.0, 100 mM NaCl). Cells were disrupted by high-pressure homogenization, and the lysate was clarified by centrifugation at 13000 rpm for 15 min at 4 ℃. The supernatant was loaded onto a Ni-NTA affinity column. The column was washed with wash buffer (25 mM Tris pH 8.0, 100 mM NaCl, 1 mM DTT, 1 mM EDTA , 15 mM imidazole), and the target protein was eluted with elution buffer (25 mM Tris pH 8.0, 100 mM NaCl, 1 mM DTT, 1 mM EDTA, 300 mM imidazole). The N-terminal His tag was removed by 3C protease digestion overnight at 4 ℃. The protein was further purified by size-exclusion chromatography on a Superdex 200 Increase column pre-equilibrated in 25 mM Tris pH 8.0, 100 mM NaCl, 1 mM DTT, 1 mM EDTA. Peak fractions corresponding to monomeric protein were collected and concentrated. Protein purity was verified by SDS-PAGE. Protein concentration was determined by NanoDrop at 280 nm . Purified protein was flash-frozen in liquid nitrogen and stored at −80 ℃.

#### Affinity Measurement between Small Molecules and MAGI2 PDZ0-GK with microscale thermophoresis

Affinity measurements between small molecules and the murine MAGI2 PDZ0– GK superdomain were performed using microscale thermophoresis (MST) on a Monolith NT.115 instrument (NanoTemper). For preliminary screening, protein concentration was fixed at 50 nM, and ligand was titrated to a maximum concentration of 25 mM. Excitation power was set automatically. Only ligands showing clear, concentration-dependent binding signals without protein aggregation were selected for quantitative affinity titration. For quantitative assays, the labeled protein was held constant, and ligand was serially diluted in MST buffer (25 mM Tris pH 8.0, 100 mM NaCl, 1 mM DTT, 1 mM EDTA) . Normalized fluorescence data were analyzed using NanoTemper Analysis software, and Kd values were obtained by nonlinear least-squares fitting assuming a 1:1 binding model based on the law of mass action. Only datasets with a signal-to-noise ratio (S/N) > 5 were used for fitting.

#### Crystal Structure Determination of the MAGI2–Ligand Complex

Purified murine MAGI2 PDZ0–GK superdomain was concentrated to 10 mg/mL. *Apo* crystals were grown at 16 ℃ using the sitting-drop vapor-diffusion method by mixing protein and reservoir solution (0.2 M MgCl₂, 0.1 M Tris-HCl pH 7.0, 28% (v/v) PEG 4000) in a 1:1 ratio. For ligand soaking, 0.5 μL of ligand stock solution was added to 0.5 μL of the drop containing mature *apo* crystals, yielding a final ligand concentration of 25 mM. The complex was incubated at 16 ℃ for at least 12 hours to allow ligand binding. Crystals were briefly soaked in cryoprotectant solution (reservoir solution supplemented with 20% (v/v) glycerol) and flash-cooled in liquid nitrogen. X-ray diffraction data were collected at 100 K at the Shanghai Synchrotron Radiation Facility (SSRF) beamlines BL18U1^61^ and BL19U1. Data were indexed, integrated, and scaled using HKL3000 or xia2-XDS. The structure was solved by molecular replacement with Phaser in PHENIX^62^ using the reported MAGI2 PDZ0– GK structure (PDB: 7YKH) as the model. Model building and manual adjustment were performed in Coot, followed by iterative refinement in PHENIX. Structural quality was validated using bond lengths, bond angles, Rwork/Rfree, and Ramachandran plots.

### Molecular dynamics simulations and trajectory analyses of MAGI2

#### Molecular dynamics simulations

Two simulation series examined the docking-derived binding mode and MAGI2 conformational dynamics. The docking-based system combined chain A of 7YKI ^41^ with the top-ranked Glide pose of MDL-6-H6; the refined complex crystal structure served as a comparison reference. The four-system series comprised *apo* MAGI2 and MAGI2–pSAPAP1 derived from 7YKI, the crystallographic MAGI2–MDL-6-H6 complex, and a ternary complex constructed by aligning MAGI2 chains and transferring pSAPAP1 from 7YKI to the MDL-6-H6-bound model.

GPU-accelerated AMBER24 simulations used ff19SB for protein ^63^, phosaa19SB for phosphorylated residues^64^ and dianionic phosphoserine in the 12-residue pSAPAP1 peptide. Systems were solvated in truncated-octahedral OPC water boxes ^65^ with a 10-Å buffer and neutralized with Na⁺ without additional salt. Following minimization, heating and restrained equilibration, production simulations used 310 K, 1 bar, Langevin temperature control (1 ps⁻¹), isotropic pressure coupling (5 ps), a 2-fs timestep, SHAKE constraints and a 10-Å direct-space cutoff. Coordinates were saved every 100 ps. The docking-based series comprised eight independent 625-ns trajectories; each of the four systems comprised four independent 1.25-μs trajectories.

#### Analysis of the docking-derived binding mode

Trajectories were reconstructed across periodic boundaries and aligned on the local pocket, applying the same transformation to protein and ligand. Analyses excluded the first 100 ns. Binding stability was assessed using crystal-referenced, symmetry-corrected ligand heavy-atom RMSD, geometric-centre displacement and residue contact occupancy. Crystallographic contact recovery used a 5.0-Å heavy-atom cutoff, whereas trajectory contacts used a 4.5-Å minimum residue–ligand heavy-atom distance. MM/GBSA and residue decomposition used 100 uniformly sampled frames per replica, MAGI2 as receptor, MDL-6-H6 as ligand, igb=5 and an implicit ionic strength of 0.15 M.

#### Interdomain geometry and shared conformational analysis

Four-system trajectories were reconstructed and aligned using matched MAGI2 residues 10–191. Primary analyses covered 200–1,250 ns at 1-ns intervals; alternative analysis windows were examined across independent replicas. Interdomain contacts joined PDZ0 residues 10–98 and GK residues 107–191 within 4.5 Å. Domain separation used non-mass-weighted Cα centroids of domain cores. After PDZ0-core alignment, the hinge angle was defined by the component of GK rigid-body rotation perpendicular to the reference interdomain axis.

Shared features comprised interface distances, pairwise distances among phosphopeptide-pocket residues and sine/cosine-transformed backbone dihedrals. Features were standardized and scaled by the square root of each block’s dimensionality, reduced to 100 principal components, and analysed using kinetic-map tICA ^66^ in deeptime. Parameters were selected by replica-held-out VAMP-2 cross-validation ^67^. The first five selected tICs were clustered using MiniBatch k-means, with model selection based on silhouette scores and bootstrap clustering stability, yielding six conformational states.

#### Phosphopeptide-binding modes

MAGI2–pSAPAP1 and ternary conformations were clustered using binary contacts between MAGI2 residues 10–191 and peptide positions −4 to +7 at a 4.5-Å heavy-atom cutoff. Contacts with pooled occupancy below 0.5% or above 99.5% were excluded. Truncated singular value decomposition and k-means identified two binding modes, assessed using Jaccard silhouette scores and bootstrap stability. The native-like mode showed the greatest Jaccard overlap with 7YKI contacts and the highest median reference-contact retention, *Q*_native_. Reference contacts were defined at 4.0 Å and their retention evaluated at 4.5 Å. The pSer–Tyr178 distance was the minimum distance between phosphate oxygen atoms and Tyr178 heavy atoms.

### Endpoint energetics and residue networks

Phosphopeptide MM/GBSA treated the complete peptide as ligand and MAGI2, together with MDL-6-H6 where present, as receptor. Overall scores used 200 post-200-ns frames per replica with igb=5 ^68^ and nested 50-frame subsets with igb=8 ^69^. Conformational entropy was omitted, and values were interpreted as endpoint interaction scores.

Replica-specific Cα cross-correlations and CorrelationPlus normalized linear mutual information ^70^ were calculated for residues 10–191 after 200 ns at 1-ns intervals. Networks connected non-adjacent residues with at least 20% contact occupancy at a 4.5-Å cutoff. Edge strength was contact occupancy multiplied by normalized mutual information, and edge cost was its negative logarithm. Shortest and suboptimal paths linked MDL-6-H6-contacting residues to the phosphopeptide-recognition region, represented by Arg143, Tyr155, Tyr178 and Tyr183. Independent replicas were the inferential units, with uncertainty estimated by replica-aware block bootstrap.

### Human proteome analysis

#### Structural data preparation and candidate pocket atlas construction

Human experimental structures from the Protein Data Bank (PDB) were integrated with predicted structures from the AlphaFold Protein Structure Database (version 4) to provide structural inputs for proteome-scale analysis. PDB structures were filtered by species, experimental method and structural quality, and small-molecule ligands, solvent and non-standard amino acid atoms were removed using Gemmi. For AlphaFold structures lacking experimental B-factors, B-factors were estimated using ResQ. PHENIX was used to remove low-confidence residues with pLDDT <70, process the estimated B values and optimize the models; the resulting B-factor estimates were used to calculate flexibility features. Structures containing no more than 1,000 amino acids were analysed using the AllositeUltra candidate pocket generation, feature extraction and scoring workflow, retaining pockets with a final composite score >0.5. To associate predictions with functional annotations, PDB residues were mapped by global sequence alignment to the UniProt human reference proteome (UP000005640) ^46^, retaining mappings with sequence identity ≥90%; AlphaFold DB predictions were associated with their corresponding UniProt entries. Predictions from both structural sources were retained. For the overall merged analysis, PDB-derived results were prioritized for each UniProt entry, with AlphaFold DB results used only when no usable PDB-derived results were available. Pocket size was defined as the number of UniProt-mapped residues in each pocket.

#### Drug-related annotations and comparison with AllositePro

Drug-related records were annotated using DGIdb and the Therapeutic Target Database (TTD).

Comparison with AllositePro was restricted to shared targets with predictions from the same structural source, following the PDB-priority rule and matching identical PDB chains. Pockets were classified as matched when the intersection of their residue sets exceeded 30% of their union or 50% of the smaller set; matched and unmatched prediction proportions were then calculated.

#### Residue composition and disease-variant enrichment

For amino acid composition analysis, residue positions from all positive pockets were pooled as a union within each protein. The relative frequencies of the 20 amino acids were compared with those in the complete sequences of the corresponding proteins, and compositional preferences were expressed as log₂ fold changes.

Disease-variant analysis used a prediction set subjected to independent mapping and sequence-completeness checks, with the complete sequences of the corresponding proteins as background. Disease-associated missense variants were obtained from UniProtKB/Swiss-Prot Humsavar ^46^, retaining likely pathogenic and pathogenic annotations. The AlphaMissense ^43^ analysis used released human missense-substitution pathogenicity predictions. Substitutions classified as pathogenic were aggregated by residue position to define four residue sets requiring at least 1, 5, 10 or 19 pathogenic substitutions per position. Pathogenic-variant frequencies, fold enrichments and odds ratios were calculated for pocket and background residues. Amino acid composition and disease-variant enrichment were assessed using Fisher’s exact tests with Benjamini–Hochberg correction for multiple comparisons.

#### Protein domains and interaction interfaces

Pfam domain annotations were obtained using PfamScan/hmmscan ^27^, retaining significant matches, and domain coverage of candidate pockets was summarized by Pfam accession. Pfam families were further grouped into clans, excluding entries without a clan assignment. Enrichment of candidate pocket residues was evaluated using one-sided hypergeometric tests against the clan-domain residues of the human proteome. Enrichment was designated at an unadjusted *P*<0.05 and fold enrichment >1, without multiple-testing correction.

Domain–domain interfaces were defined by residues making heavy-atom contacts at distances <5 Å between different Pfam domains, taking the union of contacting residues across domain pairs. Protein–protein interaction interfaces were extracted from Pinder precomputed contact maps and mapped to UniProt residue numbering. The proportions of candidate pocket and corresponding background residues overlapping each interface type were calculated, and their ratio was used to describe interface enrichment.

#### Spatial relationships between candidate pockets and functional centres

Functional centres were defined using ACT_SITE active-site annotations from the UniProtKB human reference proteome. For each candidate pocket, the geometric centroid of the Cα coordinates of all mappable pocket residues was calculated, followed by its Euclidean distance to the Cα atom of each ACT_SITE residue in the same protein:

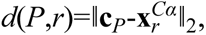

where **c***_P_* is the pocket Cα centroid and **x***^Cα^_r_* is the Cα coordinate of an active-site residue. Each pocket–active-site residue pair contributed one distance observation. Distances for experimentally annotated allosteric sites in AsBench2 were calculated using the same definition, and the two distance distributions were compared.

#### Family-level positional analysis of GPCRs and protein kinases

To compare candidate pocket positions across family members, residues in GPCR transmembrane helices TM1–TM7 were mapped to GPCRdb generic numbering ^44^. For protein kinases, catalytic domains were first located within UniProt sequences, and their residues were mapped to corresponding columns in the KinCoRe structure-based multiple sequence alignment ^45^, establishing common positional coordinates across proteins.

The highest prediction score was retained at each corresponding position within each protein, and scores >0.5 were classified as positive. Positional enrichment was assessed using binomial tests against the overall positive proportion in the corresponding family, with Benjamini–Hochberg correction for multiple comparisons. Kinase analyses additionally grouped adjacent sets of 10 alignment columns and recalculated regional enrichment and adjusted significance to display broader positional trends.

## Acknowledgements

This study was partly supported by the Innovative Drug Research and Development National Science and Technology Major Project (2025ZD1803100 & 2025ZD1803104 to J.Z.), the National Natural Science Foundation of China (82441035 to J.Z., 22237005 to J.Z.), Shanghai Municipal Health Commission (2025ZHYL038 to J.Z.), Shanghai Action Plan for Science, Technology and Innovation Field of Computational Biology (24JS2830100 to J.Z.), Lingang Laboratory (LG8888 to J.Z.), Qiankeherencai (RX [2026] 020 to J.Z.), the Natural Science Foundation of Shanghai (25ZR1401371 to W.Q.), the National Natural Science Foundation of China (82504673 to R.Z.), and Shanghai Jiao Tong University School of Medicine Graduate Innovation and Frontier Exploration (GIFE) Grant (BJ1-3000-26-1301 to Q.R.).

## Author contributions

J. Zhang conceived and supervised the project, with additional guidance from J. Zha. Q.R., X.L., J.H., W.Z. and Y.S. constructed AsBench2. Q.R. developed AllositeUltra and conducted molecular screening. Q.R., M.L. and C.W. performed computational evaluations, and Q.R. and M.L. conducted docking analyses. Q.R., M.L. and R.Z. performed molecular dynamics simulations. L.W., H.Z. and W.Q. performed binding assays and determined the crystal structures. W.F. and Z.Z. performed protein mutagenesis and activity assays. Q.R. and H.Y. conducted proteome-wide analyses. Q.R. drafted the manuscript. Q.R., J. Zhang, X.C. and R.L. discussed the results and revised the manuscript. J. Zhang assumed responsibility for the manuscript in its entirety.

## Competing interests

The authors declare no competing interests.

## Notes

### Competing Interest Statement

The authors have declared no competing interest.

